# F.A.D.E. (Fully Agentic Drug Engine): A Conversational AI Platform for Drug Discovery

**DOI:** 10.64898/2026.06.20.733481

**Authors:** Jason Kantorow, Natesan Mani, Naveen Rajagopal Mohanraj, Xue Zong

**Author notes:** These authors contributed equally to this work. Corresponding authors: Jason Kantorow, Natesan Mani.

## Abstract

Drug discovery remains one of the costliest and most time-intensive endeavors in the pharmaceutical pipeline, with average development costs exceeding $2.3 billion per drug, timelines spanning more than a decade, and attrition rates above 90% in clinical trials. While computational methods have expanded the searchable chemical space, current pipelines remain fragmented and largely inaccessible to researchers without deep interdisciplinary expertise. Here we present F.A.D.E. (Fully Agentic Drug Engine), a multi-agent, open-source platform that converts natural language queries into potential drug candidates, substantially lowering the expertise barrier to advanced computational drug discovery. F.A.D.E. employs a three-branch hierarchical architecture that adapts to the level of available structural data for any protein target, integrating structure prediction, binding pocket detection, equivariant diffusion-based de novo ligand generation, and binding affinity estimation into a single automated pipeline. We validate F.A.D.E. on two structurally distinct targets: the epidermal growth factor receptor kinase domain (EGFR), a well-established oncology target, and cellular retinol-binding protein 1 (CRBP1), a lipid-binding protein involved in retinoid metabolism. For EGFR, our generated candidates achieved QED scores of 0.85 compared to 0.46 for the co-crystallised reference ligand, demonstrating marked improvement in predicted drug-likeness. Results across both targets confirm that F.A.D.E. can reliably generate chemically tractable, drug-like hit compounds across diverse protein classes from simple natural language input.

## 1. Introduction

Drug discovery remains prohibitively expensive and time-consuming, often taking over a decade and billions of dollars to bring a single drug to market (DiMasi et al. 2016; Mullard 2014; Wouters et al. 2020). While computational approaches have emerged to accelerate this process, they face critical limitations such as proprietary constraints that lock out broader research communities, fragmented pipelines that require specialised expertise across multiple domains, and AI-driven solutions that often lack the rigorous physics-based validation necessary for reliable predictions. These challenges create significant barriers to entry and slow innovation in developing life-saving therapeutics.

There have been notable recent advances in structure-based drug design, including diffusion-based generative models (Schneuing et al. 2024; Yim et al. 2023; Wu et al. 2024), binding affinity predictors (Passaro et al. 2025), and protein structure prediction (Jumper et al. 2021; Passaro et al. 2025). However, these tools have remained largely fragmented and require considerable technical expertise, severely limiting the ability of early-stage computational researchers or experimentalists to move seamlessly from a drug discovery-based query to a viable output. Several recent efforts have sought to address this fragmentation by building LLM-powered agentic frameworks for chemical and biological research. ChemCrow (M. Bran et al. 2024) was among the first to demonstrate that augmenting an LLM with a suite of chemistry-specific tools could enable autonomous execution of drug discovery tasks. Coscientist (Boiko et al. 2023) proved that LLM agents can plan and execute chemical experiments with minimal human intervention by coordinating robotic laboratory platforms. DrugAgent (Liu et al. 2024) introduced a multi-agent framework specifically targeting the automation of machine learning programming tasks within drug discovery, achieving strong performance in ADMET prediction workflows. AgentD (Ock et al. 2026) further extended this paradigm to an end-to-end LLM-powered agent capable of data retrieval, molecular generation, multi-property prediction, and 3D protein–ligand structure generation from a single interface. Similarly, the Prompt-to-Pill (Vichentijevikj et al. 2025) framework proposes a comprehensive multi-agent architecture spanning the full preclinical pipeline, from target identification through lead optimization, synthesizing insights from over 50 LLM-based drug discovery systems. Collectively, these works establish the viability of agentic and conversational interfaces for computational drug discovery.

Despite these advances, existing agentic frameworks have largely focused on property prediction with very little to no incorporation of data obtained from robust physics-based methods. Furthermore, very few employ a data retrieval strategy that is capable of handling targets across the full spectrum of structural characterizations, from well-resolved crystal structures to sequences with no experimental structures. F.A.D.E addresses these challenges through a fully automated, open-source multi-agent workflow that aims to democratize the drug discovery process. F.A.D.E was a submission to the 2025 LLM Hackathon for Applications in Materials Science and Chemistry (Roy et al. 2026). The system accepts natural language queries, such as “Identify drug candidates targeting the EGFR receptor,” and uses these alone to orchestrate the entire pipeline from target identification and structure retrieval to de novo ligand generation, property screening, and binding affinity prediction. F.A.D.E’s novelty stems from its seamless integration of otherwise distinct tools into a cohesive, automated pipeline that handles diverse scenarios, including de novo structure prediction, equivariant diffusion-based ligand generation, binding pocket identification, and binding energy prediction. The natural language interface eliminates the need for specialised computational chemistry expertise, opening drug discovery to a wider research community. Additionally, the usage of open-source tools such as Boltz-2 (Passaro et al. 2025) and fpocket (Le Guilloux et al. 2009) expands the configurability of F.A.D.E. and provides easy access to user modification for specific needs.

Central to F.A.D.E. is a three-branch hierarchical architecture that adapts to the available structural data for any target of interest. When experimentally determined ligand-bound structures and binding site information are available in the RCSB Protein Data Bank, the system proceeds directly to de novo molecule generation (Branch 1). When a structure exists but binding site information is absent or of insufficient resolution, binding pockets are identified computationally using fpocket (Branch 2). In cases where no experimental structure exists, the target sequence is retrieved from UniProt, and the three-dimensional structure is predicted using Boltz-2, after which binding pocket identification and candidate generation proceed as in Branch 2 (Branch 3). All three branches converge on a shared candidate generation and ranking module: drug-like molecules are generated using DiffSBDD (Schneuing et al. 2024), an equivariant diffusion model conditioned on the three-dimensional binding pocket geometry, and candidates are scored and ranked by the Quantitative Estimate of Drug-likeness (QED), synthetic accessibility (SA) score, and predicted binding affinity estimated by Boltz-2.

We demonstrate the utility of F.A.D.E. on two clinically relevant protein targets: the epidermal growth factor receptor (EGFR) (PDB ID: 4G5J) (Kung 2021), a well-validated oncology target, and cellular retinol-binding protein 1 (CRBP1) (PDB ID: 5H9A) (Silvaroli et al. 2016), a less characterised target involved in retinoid metabolism. These case studies validate F.A.D.E.’s multiple-pathway architecture and illustrate its capacity to generate viable, drug-like hit compounds across diverse protein classes.

## 2. Methods

### 2.1 System Architecture and Agentic Workflow

F.A.D.E. is implemented as a multi-agentic, multi-layered workflow in which dedicated agents handle discrete stages of the drug discovery pipeline (see **Figure 1**). Users interact with the system via a natural language interface, with a dedicated agent parsing and processing the input query. Upon identifying the target of interest, this agent then directs downstream agents responsible for database retrieval, pocket identification, molecule generation, and property scoring. The modular architecture enables individual components to be updated or replaced independently as more accurate computational tools become available.

**Figure 1.**
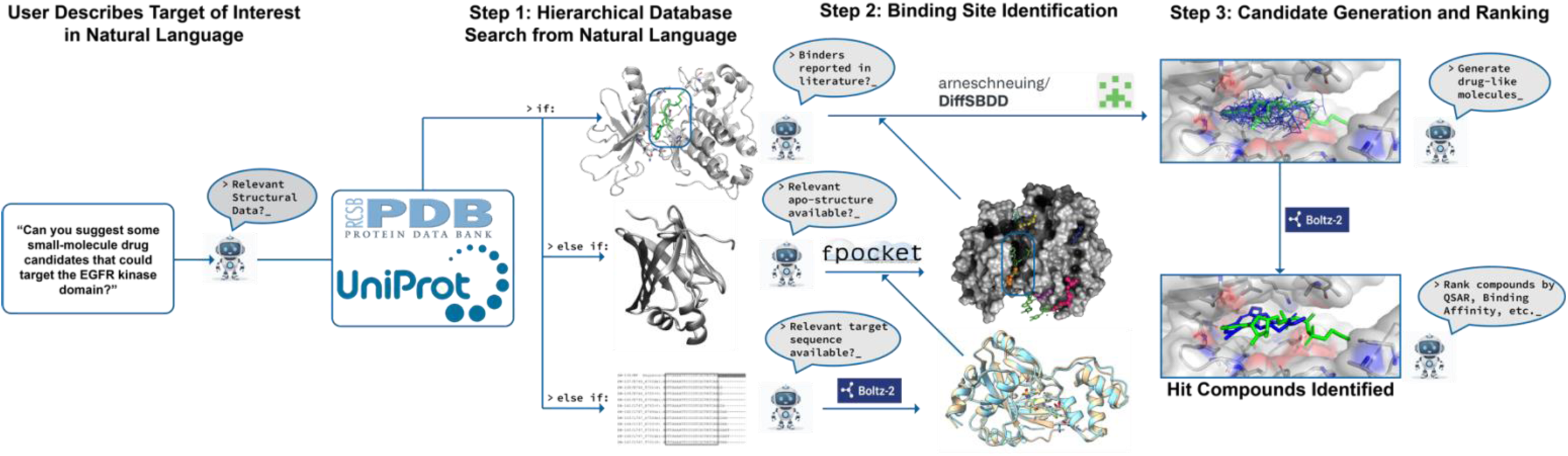
Overview of the F.A.D.E. pipeline. The workflow is initiated by a natural language query describing the target of interest. In Step 1, a hierarchical database search determines the appropriate pathway based on data availability: (i) if a ligand-bound structure with known binders is retrieved from the RCSB PDB, the pipeline proceeds directly to candidate generation; (ii) if only an apo-structure is available, binding pockets are identified computationally using fpocket; or (iii) if no experimental structure exists, the target sequence is retrieved from UniProt and the three-dimensional structure is predicted using Boltz-2. In Step 2, the identified or predicted binding site is used as input for de novo molecule generation via DiffSBDD, an equivariant diffusion model conditioned on pocket geometry. In Step 3, generated candidates are ranked by QSAR properties, Quantitative Estimate of Drug-likeness (QED), synthetic accessibility (SA) score, and predicted binding affinity estimated by Boltz-2, yielding a prioritised set of hit compounds.

### 2.2 Hierarchical Database Search and Structure Retrieval

Upon receiving a target query, F.A.D.E. conducts a hierarchical database search to identify the best structural and functional information available (see **Figure 1**). The path taken by the workflow through one of three branches depends on what data can be retrieved. Table 1 summarizes the workflow branches and their associated data requirements.

**Table 1:** Summary of F.A.D.E. workflow branches and the structural data requirements, computational tools, and confidence levels associated with each.

| Branch | Input Data | Key Tools | Pocket Source | Confidence |
| --- | --- | --- | --- | --- |
| 1 | Ligand-bound co-crystal structure | RCSB PDB | Experimental coordinates | Highest |
| 2 | Apo or unliganded structure | fpocket | Computed (fpocket) | Moderate |
| 3 | Amino acid sequence only | Boltz-2, fpocket | Computed (Boltz-2 + fpocket) | Lower |

**Branch 1: Known binders with structural data.** The agent searches the RCSB database (Burley et al. 2024) to identify an experimentally solved co-crystallised structure of the target protein in complex with known binders, including information about the specific residues or binding site coordinates with which those ligands are known to interact (see **Panel (i)** of **Figure 1**). If a relevant crystal complex between receptor and ligand(s) of sufficient resolution quality (< 3.0 Å) is found, the pipeline proceeds directly to candidate generation. The binding site is extracted by identifying receptor residues within 5Å of the present ligand, bypassing the need for computational pocket prediction. This branch represents the highest-confidence pathway, as it leverages experimentally derived structural binding data to predict novel drug discovery candidates.

**Branch 2: Structure available but binders unknown or low-resolution.** If a relevant structure is present in the PDB but known binders are unavailable, or the available co-crystallised structure is of insufficient resolution (> 3.0Å) to reliably define the binding site, F.A.D.E. then prompts an agent to perform computational binding pocket prediction using fpocket (Le Guilloux et al. 2009) (see **Panel (ii)** of **Figure 1**). fpocket is an algorithm that identifies druggable cavities on the surface of a protein and ranks them based on a druggability score. The residues of identified pockets corresponding to higher druggability scores are then used as binding site inputs for downstream de novo candidate generation.

**Branch 3: No structural data available.** If F.A.D.E is unable to find any relevant crystal structure from the RCSB database for the target of interest, the agent retrieves the protein sequence from the UniProt database (Ahmad et al. 2025) and uses Boltz-2 (Passaro et al. 2025) to predict the structure solely from its sequence (see **Panel (iii)** of **Figure 1**). Boltz-2 is a state-of-the-art, open-source, transformer-based model capable of accurate protein structure and receptor-ligand binding affinity prediction. The predicted structure is then passed to fpocket for binding pocket identification, after which the pipeline converges with **Branch 2** for candidate generation and ranking.

This hierarchical architecture ensures that F.A.D.E. can operate across the full spectrum of target characterisation, ranging from well-studied protein-ligand complexes with rich structural data to entirely uncharacterized targets for which only sequence information is available.

### 2.3 Binding Pocket Generation using fpocket

Following the hierarchical database search and structure retrieval (Section 2.2), the next critical step in the F.A.D.E. workflow is the accurate identification of the binding pockets in the protein structure. When the workflow follows **Branch 1**, the binding site is extracted directly from the experimentally determined co-crystallised ligand position, bypassing the need for computational prediction. For cases following **Branch 2** or **Branch 3**, wherein the holo structure is absent or is of low resolution, the open-source tool fpocket is used for binding pocket detection. fpocket is a robust protein cavity detection tool that identifies druggable binding sites on the protein surface by analysing the geometry of the protein and its physicochemical properties.

The algorithm identifies binding sites by calculating a druggability score for each detected pocket. This score, ranging from 0 to 1, is estimated based on several key features, including the number of alpha spheres, density, polarity, mean local hydrophobic density, and the proportion of apolar alpha spheres. The pockets that are assigned a high druggability score (typically greater than 0.5) are considered viable binding domains. These identified coordinates are then passed to the de novo ligand generation module (**Section 2.4**) to accurately determine drug-like candidates for the calculated binding pockets. This ensures that F.A.D.E. can reliably identify potential ligand-binding sites even in the absence of explicit, high-resolution experimental binding information.

### 2.4 De Novo Ligand Generation

Following binding site identification, F.A.D.E. generates drug-like small-molecule candidates using DiffSBDD (Schneuing et al. 2024). DiffSBDD is a structure-based drug design equivariant diffusion model that generates novel ligands for a particular protein pocket by incorporating translation, rotation and permutation symmetries designed around the physiochemical and topological properties of an input binding site. For each target, the top 10 generated molecules are ranked based on the Quantitative Estimate of Drug-likeness (QED) values (Bickerton et al. 2012). QED scores range from 0 to 1, with higher values indicating greater overall drug-likeness based on physicochemical properties including molecular weight, lipophilicity, hydrogen bond donors and acceptors, polar surface area, and the presence of structural alerts.

Generated molecules are filtered through a multi-metric evaluation framework prior to ranking. Only molecules meeting all acceptance thresholds simultaneously are retained for downstream binding affinity prediction. **Table 2** summarizes the evaluation criteria and thresholds applied during candidate filtering.

**Table 2:**
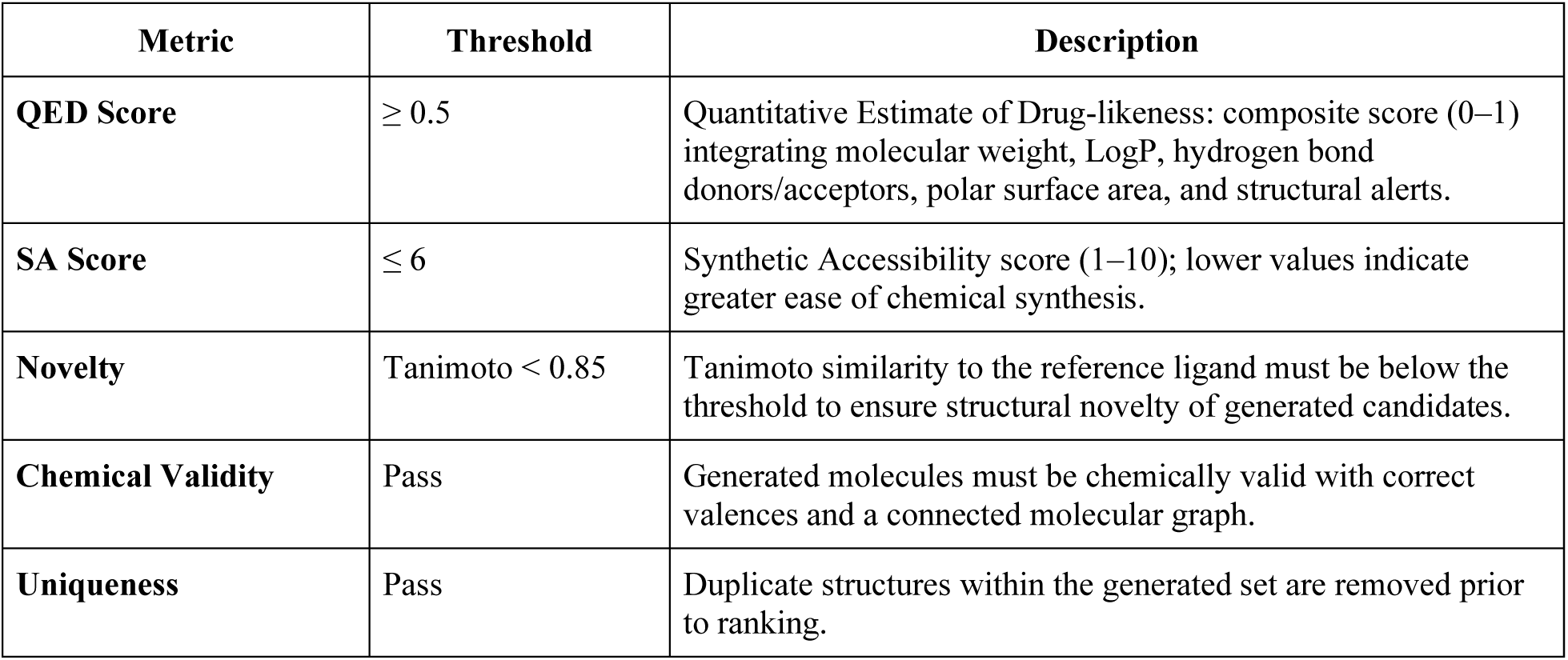
Evaluation criteria and acceptance thresholds for DiffSBDD-generated candidate molecules.

### 2.5 Binding Affinity Prediction and Candidate Ranking

Generated candidates are ranked by a combination of QED scores, synthetic accessibility (SA) score (Ertl and Schuffenhauer 2009) and predicted binding affinity values. SA scores reflect the estimated ease of chemical synthesis, with lower values indicating compounds that are more readily synthesisable. Binding affinities are estimated using Boltz-2, which provides rapid binding affinity predictions from protein-ligand complexes without relying on molecular dynamics simulations or rigorous free energy calculations. This integration of these metrics allows F.A.D.E. to prioritize candidates that are not only geometrically complementary to the target pocket but also drug-like, synthesizable, and predicted to be high-affinity binders.

### 2.6 Case Study Targets

F.A.D.E. was validated on two targets. The first, EGFR, is a receptor tyrosine kinase that is well-established in lung cancer-targeting therapies. Both apo and holo structures were readily extracted from the RCSB database, making this target well-suited to demonstrate Branch 1 of the workflow (see **section 2.2** in **Methods**). The second, CRBP1, is a lipid-binding protein involved in vitamin A (retinol) transport and metabolism. While both apo and holo conformations were found in the RCSB database, the holo structures are generally of lower resolution. This target was therefore selected to demonstrate Branch 2 of the workflow.

**Branch 3** of the F.A.D.E. workflow is designed to perform de novo structure prediction from sequence via Boltz-2, followed by fpocket-based pocket identification and DiffSBDD-based ligand generation. In practice, the majority of commonly studied drug targets have at least an apo structure deposited in the PDB, so **Branches 1** and **2** cover most routine use cases. **Branch 3** is intended primarily for novel or poorly characterized targets for which no experimental structure exists. While the platform’s architecture supports this pathway, its full implementation has not yet been completed, and the two case studies presented here exercise **Branches 1** and **2** only. Completing and validating **Branch 3**, including benchmarking the predicted structures against experimentally determined ground-truth structures, remains a priority for extending F.A.D.E.’s coverage to this harder regime and is an immediate next step for this work.

## 3. Results

### 3.1 Validation of Binding Pocket Identification

A critical design decision in the F.A.D.E. pipeline was the selection of fpocket as the binding pocket detection algorithm for **Branches 2** and **3** of the workflows. To validate this choice, we assessed fpocket’s ability to recover experimentally known binding sites from an apo structure by comparing its predictions against a co-crystallised holo structure of the same target.

We obtained the apo structure of EGFR (PDB: 7SI1) and evaluated fpocket’s predictions by comparison to the experimentally determined holo structure (PDB: 4G5J), which contains two co-crystallised ligands: 0WN, bound in the ATP-binding cleft, and 0WM. fpocket’s highest-scoring pocket on the apo structure, Pocket 15 (druggability score = 0.713), overlaps precisely with the experimentally characterised 0WN binding site (see SI Figure 1). The secondary pockets, Pocket 9 (druggability = 0.154) and Pocket 4 (druggability = 0.139), overlap with the 0WM ligand positions. These results confirm that fpocket can reliably recover known binding sites from apo structures, validating its incorporation as the pocket detection tool in the F.A.D.E. workflow.

### 3.2 Pipeline Performance on EGFR

For the EGFR target, F.A.D.E. successfully retrieved the holo crystal structure from the RCSB database, with two co-crystallised ligands 0WN and 0WM. F.A.D.E identified the binding sites using these ligands as a reference following the **Branch 1** pathway. A diverse set of small-molecule drug candidates was then generated for each of these binding sites using DiffSBDD. The top 10 candidates for each binding site were then ranked by QED and SA scores and considered for binding affinity evaluation using Boltz-2. The results were then compared against reference ligand poses.

Both the reference docked ligands 0WN and 0WM exhibited a QED score of 0.46 and an SA score of 3.08, with a Boltz-2 predicted binding affinity of –1.31 (**Table 3**). Based on the binding site obtained from 0WN, the top two candidates generated by F.A.D.E (Ligand 1 and Ligand 2) exhibited substantially improved QED scores of 0.85, indicating better predicted drug-likeness compared to the reference bound ligand. The SA scores of these two candidates were 4.51 and 3.99, respectively, suggesting moderate ability to synthesize. The predicted binding affinities (–0.32 and –0.23 respectively) were higher (weaker) than the reference ligand 0WN, a finding that is discussed further in Section 4. In the case of the binding site obtained from 0WM, the predicted affinity values indicate weak binders or almost non-binders. It is important to note that the generated ligands are novel drug candidates selected to balance multiple desired properties and are not optimized for binding affinity alone. The primary objective of this initial pipeline is the identification of drug-like scaffolds for further optimization and hence downstream methods are planned to increase the binding affinity of chosen candidates.

**Table 3:**
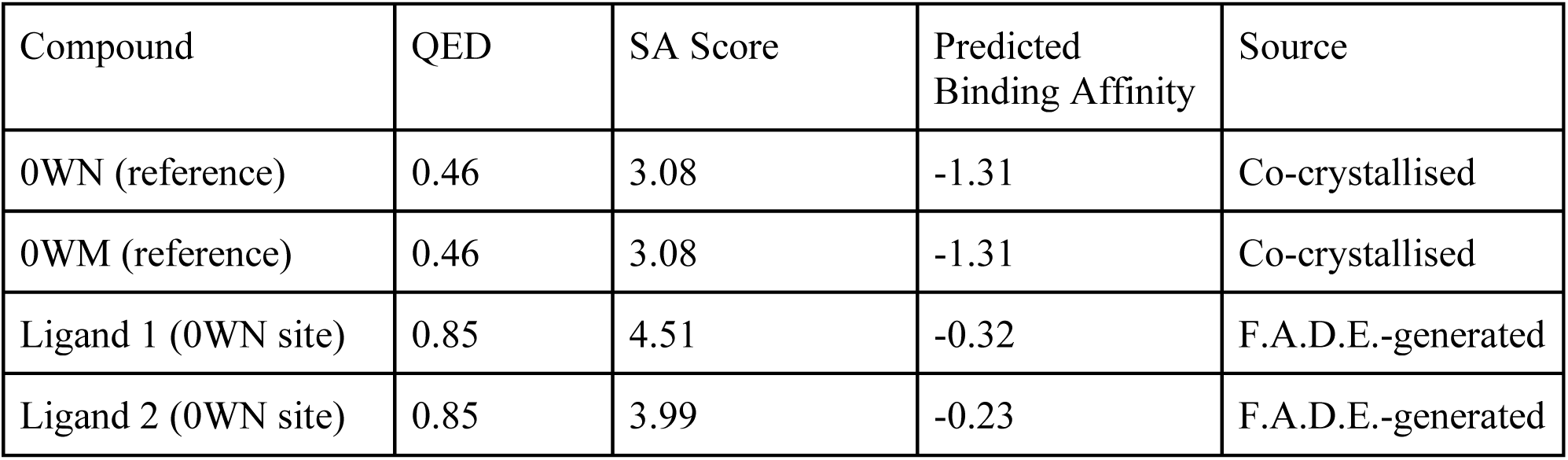
Comparison of QED, SA, and predicted binding affinity scores for EGFR reference 246 ligands and top F.A.D.E. generated candidates.

The chemical space coverage of generated EGFR candidates was visualised (see **Panel C** of **Figure 2**) using t-SNE dimensionality reduction (van der Maaten 2008), confirming that F.A.D.E. generates structurally diverse molecules spanning distinct regions of chemical space relative to the reference ligand. Candidates were generated from three independent binding pockets identified by the pipeline, further expanding structural diversity.

**Figure 2.**
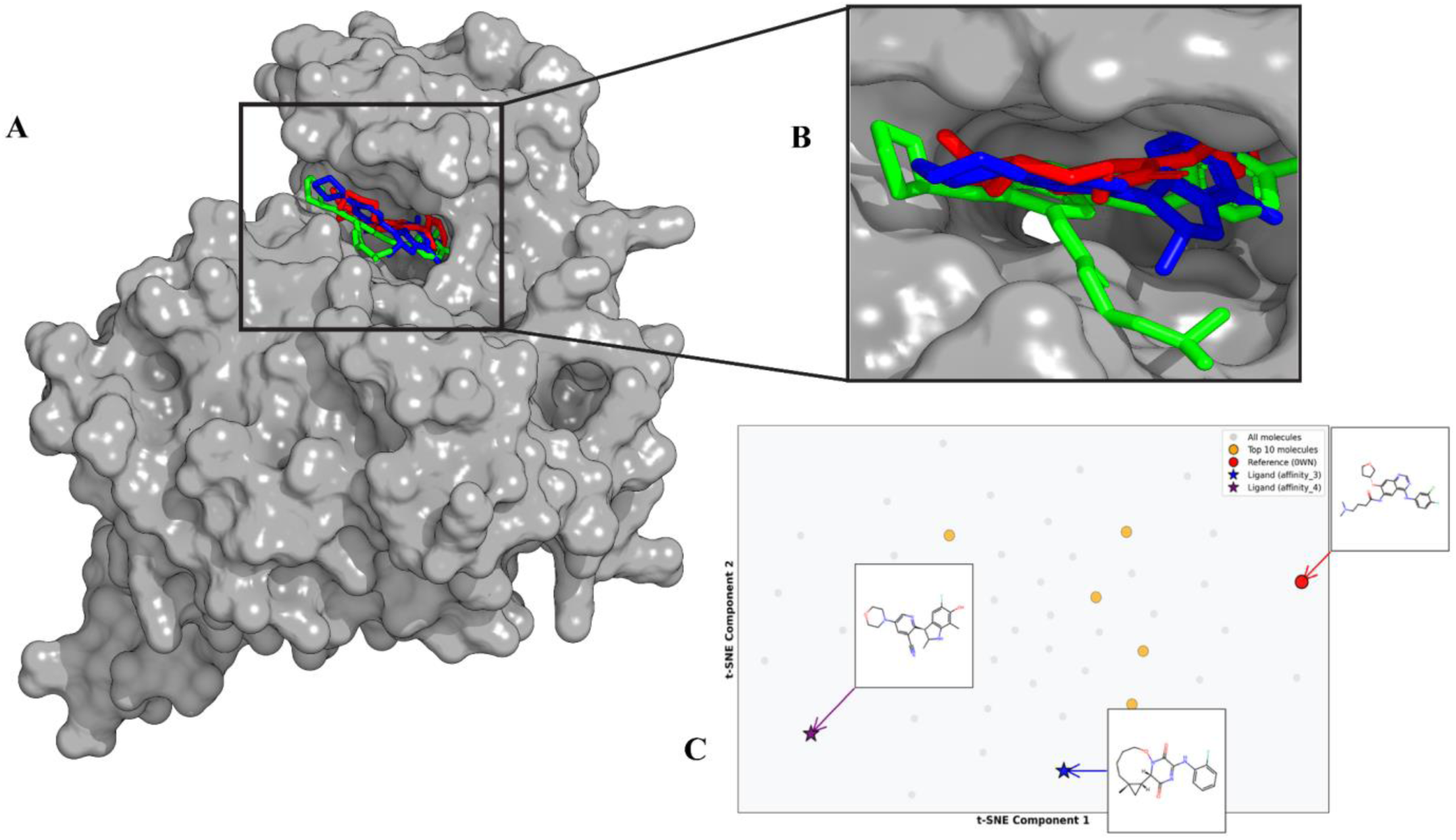
F.A.D.E.-generated small-molecule candidates docked to the EGFR kinase domain. Surface representation of the EGFR kinase domain, with the ATP-binding pocket highlighted by the inset region (see **Figure 2A**). In **Figure 2B**, we see a close-up view of the binding pocket showing the co-crystallised reference ligand (0WN, shown in red) overlaid with the two top-ranked F.A.D.E.-generated candidates, named affinity_3 (shown in green) and affinity_4 (shown in blue), demonstrating geometrically plausible binding poses within the pocket. In **Figure 2C**, we see t-SNE dimensionality reduction of the full generated library of compounds (shown in grey), with the top 10 QED-ranked molecules (highlighted in orange), the reference ligand 0WN (red circle), and the two top affinity-ranked candidates, affinity_3 (purple star) and affinity_4 (blue star), annotated with their two-dimensional chemical structures. The spatial separation of generated candidates from the reference ligand confirms that F.A.D.E. samples a chemically diverse region of drug-like chemical space.

### 3.3 Pipeline Performance on CRBP1

In the case of the CRBP1 target, F.A.D.E correctly selected **Branch 2** of the workflow for pocket detection, as a high-resolution holo structure was not available. fpocket was used to determine binding pockets based on the apo structure from RCSB and identified nine potential binding pockets on the protein surface. Table 4 summarises the key physicochemical and geometric properties of all detected pockets. Only four pockets exhibited non-zero druggability scores (Pockets 1, 5, 6, and 8), with Pocket 1 showing a substantially higher druggability score (0.888) than all others.

**Table 4:** Physicochemical properties of all binding pockets identified by fpocket analysis of CRBP1 (PDB: 5H9A). Pockets are listed in order of pocket number as assigned by fpocket.

| Pocket | Score | Druggability | Volume (Å<sup>3</sup>) | SASA | Apolar Ratio | Hydrophobicity | Alpha Spheres |
| --- | --- | --- | --- | --- | --- | --- | --- |
| 1 | 0.350 | 0.888 | 668.4 | 175.0 | 0.585 | 40.3 | 106 |
| 2 | 0.060 | 0.000 | 382.8 | 85.6 | 0.188 | 11.6 | 16 |
| 3 | 0.051 | 0.000 | 500.3 | 156.2 | 0.206 | 9.8 | 34 |
| 4 | 0.043 | 0.000 | 170.4 | 23.1 | 0.474 | 14.4 | 19 |
| 5 | 0.027 | 0.001 | 395.4 | 105.7 | 0.296 | 28.6 | 27 |
| 6 | -0.004 | 0.004 | 288.8 | 105.5 | 0.560 | 23.1 | 25 |
| 7 | -0.026 | 0.000 | 223.3 | 74.8 | 0.267 | 12.8 | 15 |
| 8 | -0.063 | 0.002 | 289.8 | 118.4 | 0.600 | 37.7 | 15 |
| 9 | -0.064 | 0.000 | 228.7 | 82.8 | 0.150 | 24.8 | 20 |

Pocket 1 is characterised by a large volume (668.4 Å³), a high apolar alpha-sphere proportion (0.585), and a hydrophobicity score of 40.3, consistent with the deeply buried hydrophobic channel that naturally accommodates retinol in this protein family. The remaining druggable pockets (5, 6, and 8) exhibited markedly lower druggability scores (≤ 0.004), representing potential secondary or allosteric binding sites. DiffSBDD was run independently for each druggable pocket, and the top 10 candidates by QED across all pockets originated from Pockets 1, 5, and 6.

The most promising candidates, based on a combination of QED score and predicted binding characteristics from Boltz-2, were identified and named as Molecule_03_Pocket_1, Molecule_06_Pocket_6, and Molecule_08_Pocket_5. This finding indicates that viable candidates can be generated across multiple identified binding sites even when high-resolution holo-structure data is unavailable, confirming the utility of Branch 2. Visual inspection of the generated candidates in the binding pocket confirms geometrically possible binding poses for multiple top-ranked compounds (see **Panels B** and **C** of **Figure 3).**

**Figure 3.**
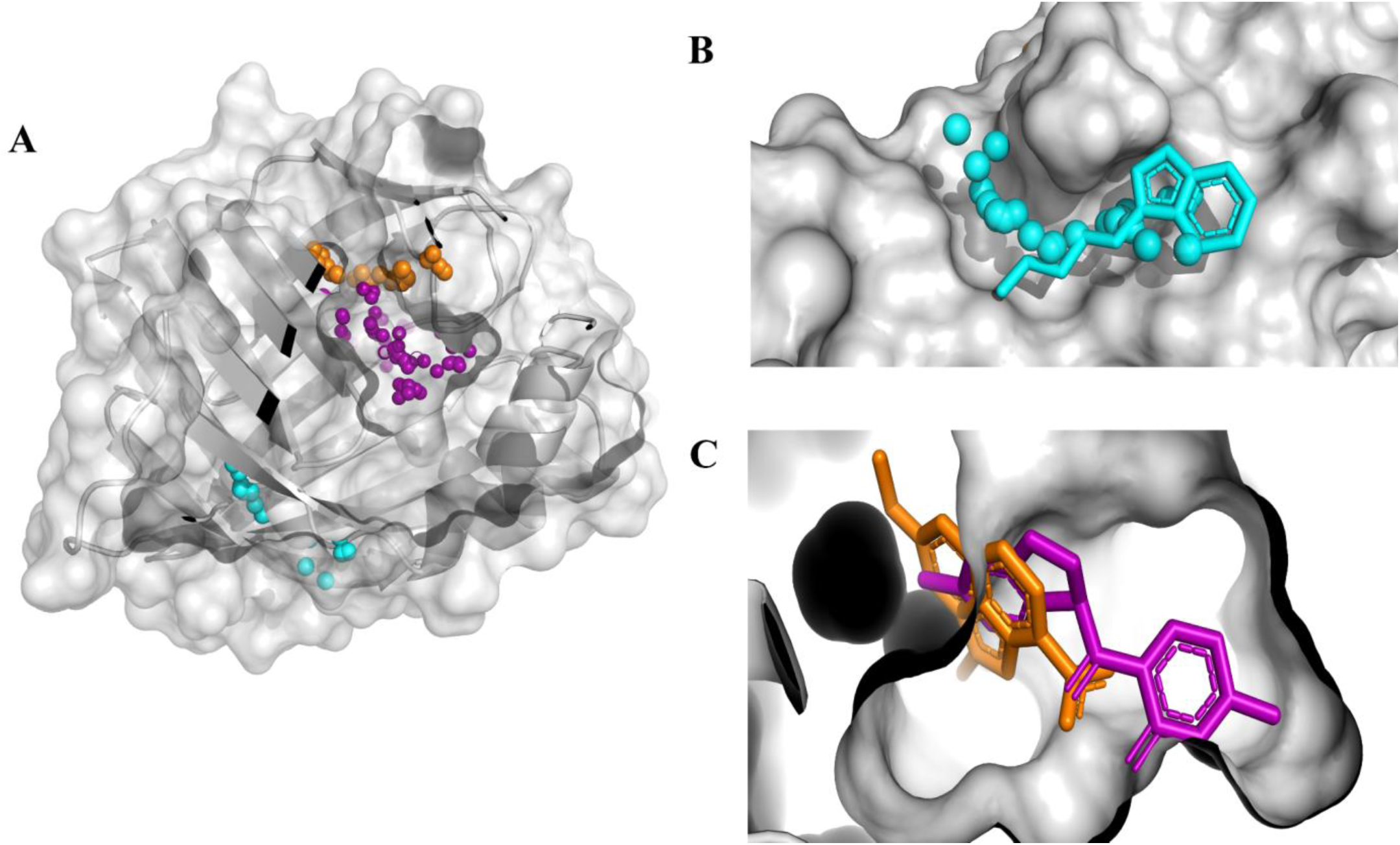
F.A.D.E.-generated small-molecule candidates docked to CRBP1. Combined surface and ribbon representation of CRBP1 are shown in **Figure 3A**, showing the distribution of generated candidates across three distinct binding pockets identified by the pipeline. We see the representations of the primary retinol-binding cavity (shown in magenta), a secondary surface-exposed pocket (shown in orange), and a third peripheral site (shown in cyan).In **Figure 3B**, we see a close-up view of the cyan pocket showing a top-ranked candidate in stick representation, occupying a shallow surface groove on the protein exterior. In **Figure 3C**, we see a close-up view of the primary binding cavity showing the two top-ranked candidates, one from the orange pocket and one from the magenta pocket. They both adopting geometrically complementary poses within the deep hydrophobic interior characteristic of lipid-binding proteins. The multi-pocket sampling demonstrated here illustrates F.A.D.E.’s ability to explore diverse binding sites on a single target, expanding the hit compound search space beyond a single predefined cavity.

## 4. Discussion

Our findings demonstrate that F.A.D.E. represents a meaningful step toward the democratization and large-scale integration of computational drug discovery tools. By combining natural language interaction with agentic AI, cutting-edge physics-based tools, and binding affinity predictions within a single automated workflow, the platform removes key barriers that have confined advanced computational methods to specialised research groups. The results on EGFR and CRBP1 demonstrate that F.A.D.E. can reliably generate drug-like candidates across chemically diverse targets, providing a foundation for broader therapeutic exploration.

Compared to existing agentic frameworks, F.A.D.E. offers several distinguishing features. Unlike ChemCrow (M. Bran et al. 2024) and DrugAgent (Liu et al. 2024), which primarily focus on tool augmentation for property prediction and ADMET workflows, F.A.D.E. integrates structure-based de novo ligand generation through DiffSBDD, producing entirely novel molecular scaffolds rather than screening existing compound libraries. AgentD (Ock et al. 2026) shares F.A.D.E.’s end-to-end ambition but relies on a single linear pipeline, whereas F.A.D.E.’s three-branch hierarchical architecture allows the system to adapt its strategy to the level of available structural data. The Prompt-to-Pill framework (Vichentijevikj et al. 2025) proposes the broadest scope, spanning the full preclinical pipeline, but remains largely conceptual; by contrast, F.A.D.E. presents validated results on multiple protein targets. Additionally, F.A.D.E.’s reliance on open-source, physics-based tools such as Boltz-2 and fpocket distinguishes it from frameworks that depend on proprietary models or purely data-driven property predictors without structural grounding.

From the EGFR case study, the substantial improvement in QED scores observed for FADE-generated candidates (0.85 versus 0.46 for the reference co-crystallised ligand) suggests that the generative pipeline inherently favours drug-like chemical compounds. This is consistent with the design of DiffSBDD, which samples from a learned distribution over drug-like chemical space conditioned on pocket geometry, rather than optimising against a single binding affinity objective. While the predicted binding affinities of generated compounds did not surpass the reference ligand, this outcome reflects the intended design of the current pipeline, which prioritises diverse, drug-like scaffold generation over single-property optimization.

The binding affinity results warrant careful interpretation. The weaker predicted affinities of F.A.D.E.-generated candidates relative to the co-crystallised reference ligands do not necessarily indicate poor therapeutic potential, since the reference ligands have undergone extensive medicinal chemistry optimisation, whereas the generated candidates represent initial hit compounds. It is also important to note that Boltz-2’s affinity estimation is not a simple scoring function applied to a rigid receptor: the model jointly predicts the three-dimensional arrangement of both the protein and ligand in the bound state, meaning the receptor conformation in the output may differ from the input crystal structure as the model predicts conformational adjustments upon ligand binding. The affinity values are therefore derived from these predicted complexes rather than from rigid docking. Nonetheless, these predictions have not been systematically benchmarked against experimental binding data for de novo generated compounds, and the absolute values should be interpreted with caution. The primary value of these predictions within the current pipeline lies in their use as a relative ranking metric among generated candidates rather than as absolute measures of binding strength. Future iterations of F.A.D.E. will incorporate more rigorous affinity estimation methods, such as molecular dynamics-based free energy perturbation or MM-GBSA rescoring, to provide higher-confidence affinity predictions and enable direct comparison with experimental IC50 or Kd values.

The modular architecture of F.A.D.E. is a significant design strength. The individual modules—structure retrieval, pocket detection, molecule generation, and affinity prediction—function independently, enabling users to substitute bespoke or cutting-edge tools as they become available. This flexibility positions F.A.D.E. as a living platform capable of incorporating methodological advances in real time.

Several limitations of the current implementation should be acknowledged. The SA scores indicate that some generated candidates may present synthetic challenges, with scores above 4.0 suggesting moderately complex synthesis routes. Addressing these limitations through integration of retrosynthetic planning tools and expanded experimental validation is a priority for future development.

## 5. Conclusion

We have introduced F.A.D.E., a fully agentic, open-source drug discovery platform that enables researchers to generate and evaluate small-molecule drug candidates through natural language queries. By orchestrating a hierarchical workflow that adapts to the level of available structural data, F.A.D.E. demonstrates robust performance across distinct protein targets.

The platform’s key achievements include widening the accessibility of computational drug discovery tools through a conversational interface, a multiple-pathway architecture that maintains performance regardless of whether experimental structural data is available, and the generation of drug-like candidates with QED scores substantially exceeding those of reference ligands. Future development will focus on systematic benchmarking against experimental binding data, incorporation of more rigorous binding affinity estimation methods, expansion to additional therapeutic modalities, automated candidate docking for additional binder screening, and broader validation across diverse target classes.

F.A.D.E. is openly available on GitHub (https://github.com/Naveen-R-M/F.A.D.E), and we invite the community to contribute to its continued development.

## Author Contributions

N.M.: Conceptualisation, Data Curation, Writing, Analysis, Visualisation; J.K.: Software, Data Curation, Analysis, Visualisation; N.R.M.: Software, Data Curation, Analysis; X.Z.: Conceptualisation, Software, Data Curation and Analysis, Writing.

## Notes

### Competing Interest Statement

The authors have declared no competing interest.

## References

1. Ahmad S, da Costa Gonzales LJ, Bowler-Barnett EH, Rice DL, Kim M, Wijerathne S, Luciani A, Kandasaamy S, Luo J, Watkins X, Turner E, Martin MJ, the UniProt Consortium. 2025. The UniProt website API: facilitating programmatic access to protein knowledge. Nucleic Acids Research. 53(W1):W547–W553. doi:10.1093/nar/gkaf394.

2. Bickerton GR, Paolini GV, Besnard J, Muresan S, Hopkins AL. 2012. Quantifying the chemical beauty of drugs. Nature Chemistry. 4(2):90–98. doi:10.1038/nchem.1243.

3. Boiko DA, MacKnight R, Kline B, Gomes G. 2023. Autonomous chemical research with large language models. Nature. 624(7992):570–578. doi:10.1038/s41586-023-06792-0.

4. Burley SK et al. 2024. Updated resources for exploring experimentally-determined PDB structures and computed structure models at the RCSB Protein Data Bank. Nucleic Acids Research. 53(D1):D564–D574. doi:10.1093/nar/gkae1091.

5. DiMasi JA, Grabowski HG, Hansen RW. 2016. Innovation in the pharmaceutical industry: new estimates of R&D costs. Journal of Health Economics. 47:20–33. doi: 10.1016/j.jhealeco.2016.01.012.

6. Ertl P, Schuffenhauer A. 2009. Estimation of synthetic accessibility score of drug-like molecules based on molecular complexity and fragment contributions. Journal of Cheminformatics. 1(1):8. doi:10.1186/1758-2946-1-8.

7. Jumper J et al. 2021. Highly accurate protein structure prediction with AlphaFold. Nature. 596(7873):583–589. doi:10.1038/s41586-021-03819-2.

8. Kung JE, Wu P, Kiefer JR, Sudhamsu J. 2021. Crystal structure of apo EGFR kinase domain. Protein Data Bank. PDB ID: 7SI1.

9. Le Guilloux V, Schmidtke P, Tuffery P. 2009. Fpocket: an open source platform for ligand pocket detection. BMC Bioinformatics. 10:168. doi:10.1186/1471-2105-10-168.

10. Liu S, Lu Y, Chen S, Hu X, Zhao J, Lu Y, Zhao Y. 2024. DrugAgent: automating AI-aided drug discovery programming through LLM multi-agent collaboration. arXiv. arXiv:2411.15692.

11. M. Bran A et al. 2024. Augmenting large language models with chemistry tools. Nature Machine Intelligence. 6(5):525–535. doi:10.1038/s42256-024-00832-8.

12. Mullard A. 2014. New drugs cost US$2.6 billion to develop. Nature Reviews Drug Discovery. 13(12):877. doi:10.1038/nrd4507.

13. Ock J et al. 2026. Large language model agent for modular task execution in drug discovery. Journal of Chemical Information and Modeling. 66(4):2055–2068. doi:10.1021/acs.jcim.5c02454.

14. Passaro S, Corso G, Wohlwend J, Reveiz M, Thaler S, Somnath VR, Getz N, Portnoi T, Roy J, Stark H, Kwabi-Addo D, Beaini D, Jaakkola T, Barzilay R. 2025. Boltz-2: towards accurate and efficient binding affinity prediction. bioRxiv. doi:10.1101/2025.06.14.659707.

15. Roy A et al. 2026. From knowledge to action: outcomes of the 2025 large language model (LLM) hackathon for applications in materials science and chemistry. arXiv. arXiv:2605.03205.

16. Schneuing A, Harris C, Du Y, Didi K, Jamasb A, Igashov I, Du W, Gomes C, Blundell TL, Lio P, Welling M, Bronstein M, Correia B. 2024. Structure-based drug design with equivariant diffusion models. Nature Computational Science. 4(12):899–909. doi:10.1038/s43588-024-00737-x.

17. Silvaroli JA, Arne JM, Chelstowska S, Kiser PD, Banerjee S, Golczak M. 2016. Ligand binding induces conformational changes in human cellular retinol-binding protein 1 (CRBP1) revealed by atomic resolution crystal structures. Journal of Biological Chemistry. 291(16):8528–8540. doi:10.1074/jbc.M116.714535.

18. van der Maaten L, Hinton G. 2008. Visualizing data using t-SNE. Journal of Machine Learning Research. 9:2579–2605.

19. Vichentijevikj I, Mishev K, Simjanoska Misheva M. 2025. Prompt-to-pill: multi-agent drug discovery and clinical simulation pipeline. Bioinformatics Advances. 6(1). doi:10.1093/bioadv/vbaf323.

20. Wouters OJ, McKee M, Luyten J. 2020. Estimated research and development investment needed to bring a new medicine to market, 2009-2018. JAMA. 323(9):844–853. doi:10.1001/jama.2020.1166.

21. Wu KE et al. 2024. Protein structure generation via folding diffusion. Nature Communications. 15(1):1059. doi:10.1038/s41467-024-45051-2.

22. Yim J, Stärk H, Corso G, Jing B, Barzilay R, Jaakkola TS. 2024. Diffusion models in protein structure and docking. WIREs Computational Molecular Science. 14(2):e1711. doi:10.1002/wcms.1711.

