## Supplemental Information for "F.A.D.E. (Fully Agentic Drug Engine): A Conversational AI Platform for Drug Discovery"

### S1. Agentic Workflow Adaptive Behaviour

A key architectural feature of F.A.D.E. is its agentic design, in which each pipeline stage is managed by an independent agent capable of autonomous decision-making and adaptive behaviour based on intermediate results. Table S1 summarises the workflow branches and their associated data requirements. This section describes the primary adaptive mechanisms implemented in the current system.

#### S1.1. Adaptive Batch Size Adjustment

The molecule generation agent monitors the pass rate of generated candidates against the evaluation criteria defined in Table 2 of the main manuscript. If the initial batch of 50 molecules yields fewer than 25 passing candidates (below the 50% target pass rate), the agent automatically increases the batch size for subsequent generation rounds. This ensures that a sufficient number of high-quality candidates are produced even for binding pockets with challenging geometries, without requiring manual intervention. Table S1 summarizes the adaptive generation parameters.

| Parameter | Value |
| --- | --- |
| **Initial batch size** | 50 molecules |
| **Maximum batches** | 3 |
| **Target passing candidates** | 25 (50% pass rate) |
| **Batch size adjustment** | Automatic increase based on observed pass rate |
| **Final ranking metric** | QED score (top 10 selected for Boltz-2 evaluation) |

**Table S1.** Adaptive molecule generation parameters used by the F.A.D.E. agentic workflow.

#### S1.2. Multi-Pocket Merging Strategy

When fpocket identifies multiple druggable pockets on a target protein, DiffSBDD is run independently for each pocket using the pocket-specific binding residues as spatial conditioning. The agent then merges results from all productive pockets into a single ranked list ordered by QED score before selecting the top 10 candidates for Boltz-2 binding affinity prediction. For CRBP1, this strategy resulted in candidates from three distinct pockets (Pockets 1, 5, and 6) appearing in the final top 10, maximizing the structural diversity of the candidate set.

### S2. Validating fpocket as the Pocket Detection Method

SI Figure 1 provides a visual representation of all 19 fpocket-identified binding pockets on the EGFR kinase domain surface. This visualization supplements the quantitative validation presented in Section 3.1 of the main manuscript.

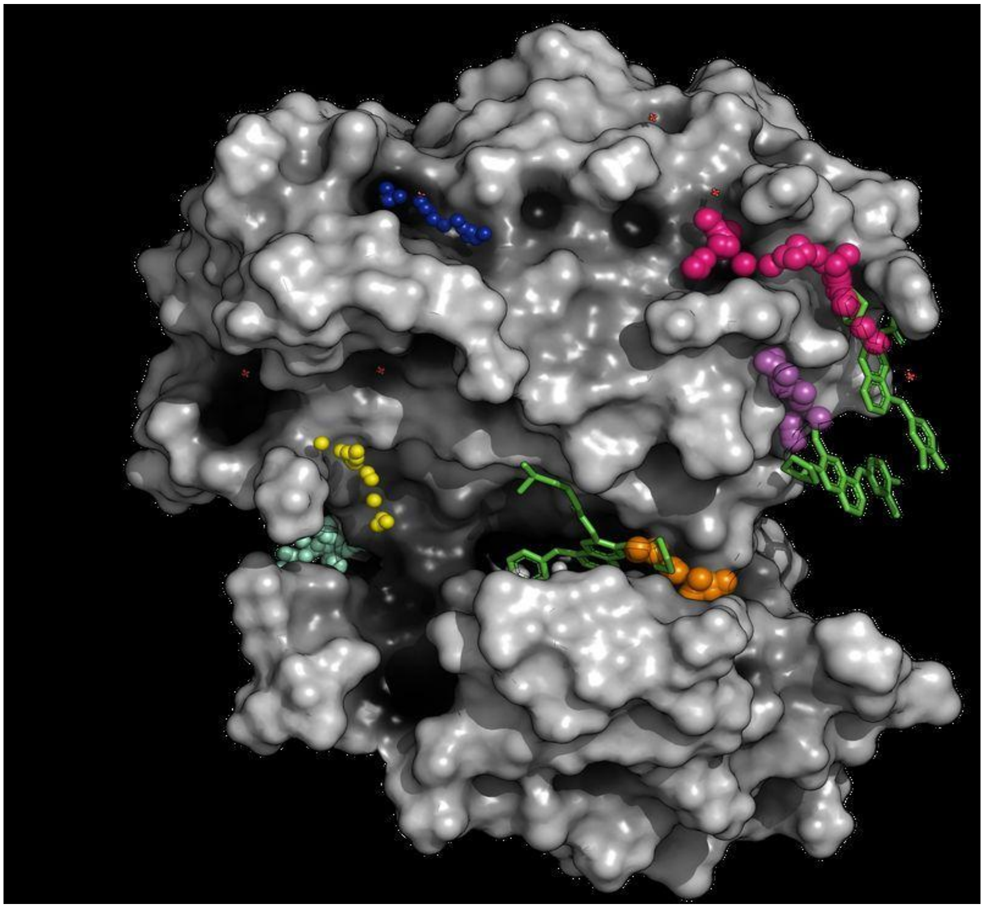

**SI Figure 1:** Surface representation of the EGFR kinase domain (PDB id: 4G5J) with all 19 fpocket-identified binding pockets visualized as colored alpha-sphere clusters. Each color represents a distinct pocket. The largest cluster corresponds to Pocket 15 (see pocket corresponding to orange spheres), which overlaps with the experimentally determined ATP-binding cleft containing the co-crystallised reference ligand 0WN. We also observe Pockets 4 and 9 overlapping with the co-crystallized 0WM ligands (shown in magenta and purple spheres, respectively). The accurate identification of the ligand binding sites by fpocket validates the reliability of this tool and its incorporation in the workflow.

### S3. Boltz-2: Complex Structure Prediction and Binding Affinity Estimation

Boltz-2 serves two distinct roles within the F.A.D.E. pipeline, and it is important to clarify the nature of its contribution at each stage. In Branch 3 of the workflow (when no experimental structure is available), Boltz-2 performs de novo protein structure prediction from the amino acid sequence retrieved from UniProt, generating a three-dimensional apo structure for downstream pocket detection and ligand generation. In this role, Boltz-2 functions as a structural prediction engine similar to AlphaFold.

Across all three branches, Boltz-2 is subsequently used for binding affinity estimation of the top 10 DiffSBDD-generated candidates. A condensed description of this dual role and its implications for affinity interpretation is provided in Section 4 of the main manuscript; this section provides additional methodological detail.

For the EGFR case study (PDB ID: 4G5J), the input receptor structure was the experimentally determined holo crystal structure. Boltz-2 re-predicted the protein–ligand complex for each of the top 10 candidates, and the resulting receptor conformations showed minor deviations from the input crystal structure, primarily in flexible loop regions distal to the binding site. The core binding pocket geometry remained largely preserved, suggesting that for well-resolved crystal structures, Boltz-2's complex prediction does not substantially alter the receptor conformation. For the CRBP1 case study, where the input structure was the apo form (PDB: 5H9A), the potential for Boltz-2 to predict meaningful binding-induced conformational changes is greater, though systematic comparison of receptor RMSD values between input and output structures was not performed in this work and represents a direction for future validation.

The binding affinity predictions from Boltz-2 should be interpreted as relative rankings rather than absolute free energy values. As noted in the manuscript, the predicted affinity values for both EGFR and CRBP1 candidates indicated weak to moderate predicted binding, which is consistent with the expected performance of de novo generative models at the hit-identification stage. One practical limitation observed was a conformer generation failure for one CRBP1 candidate (Molecule 5 from Pocket 5), indicating that certain molecular geometries produced by DiffSBDD may fall outside the conformer sampling range of Boltz-2. This represents an area for future pipeline robustness improvement.
